# Pediatric CHOP Chemotherapy Acutely Disrupts Satellite-Cell Dynamics and Blunts Muscle Mass in a Sex-Specific Manner

**DOI:** 10.1101/2025.08.04.668538

**Authors:** Jainil Daredia, Marc A Magaña, Carla MC Nascimento, Jaden M Wells, Nicholas T Thomas, Yuan Wen, Savannah V Rauschendorfer, Cory M Dungan, Michael P Wiggs

## Abstract

Pediatric cancer survival now exceeds 85 percent owing, in part, advances and use of combination chemotherapy treatments such as CHOP (cyclophosphamide, doxorubicin, vincristine, and prednisone). Despite its efficacy, CHOP may cause deleterious off-target effects during critical pediatric development periods such as impairments of skeletal muscle. We evaluated the acute effects of a CHOP administered to C57Bl/6J mice from postnatal day 28 to 48. CHOP slowed body-weight gain and has smaller gastrocnemius fiber cross-sectional area by approximately 25 percent in both sexes. mRNA sequencing detected 214 differentially expressed genes in males and 217 in females relative to controls, yet only 29 transcripts overlapped. Males exhibited downregulation of myogenic regulators, indicating impaired progenitor maintenance, whereas females showed an upregulation of extracellular-matrix and translational machinery genes plus cell-cycle regulators. Using immunohistochemistry to assess satellite cell abundance, there were 60% fewer satellite cells in males and a 40% fewer in females, which supported our transcriptional findings. These results demonstrate that pediatric CHOP acutely disrupts muscle stem-cell dynamics via sex-specific molecular programs and identify satellite-cell depletion as a potential target for preserving muscle health in pediatric cancer survivors.

## Introduction

Pediatric cancer survival has improved dramatically, with over 85% of childhood, juvenile, and adolescent and young adult (AYA) patients now living at least five years post-diagnosis (1). The improvement in survivorship stems largely from optimized chemotherapy regimens, yet systemic side effects persist. Chemotherapy-associated blunting of skeletal muscle growth and development poses a particular threat during adolescence, a critical period for skeletal muscle growth that underpins long-term metabolic health and functional capacity (2).

CHOP, a multi-agent regimen consisting of <u>c</u>yclophosphamide, <u>h</u>ydroxydaunorubicin (doxorubicin), <u>o</u>ncovin (vincristine), and <u>p</u>rednisone, forms the backbone of pediatric non-Hodgkin lymphoma treatment. Moreover, each of the single chemotherapeutic agents are often used for treatment of a variety of pediatric malignancies (3). Individually, components of CHOP impair muscle homeostasis: cyclophosphamide reduces gastrocnemius mass in rodents (4), doxorubicin triggers oxidative stress and mitochondrial dysfunction (5, 6), vincristine induces peripheral neuropathy (7), and glucocorticoids exacerbate atrophy (8). When combined, this collection of clinically relevant agents likely accelerates muscle decline and further disrupts muscle development.

Despite these well-documented single-agent effects of some chemotherapies, preclinical pediatric studies of multi-agent regimens, which combine drugs with distinct mechanisms and non-overlapping toxicities, remain scarce (9, 10) even though such combination therapies are the clinical standard. This gap is critical in juvenile models, where growing muscles undergo both fiber hypertrophy and longitudinal growth, which require coordinated processes such as activation of satellite cells for myonuclear accretion, enhancement of ribosomal biogenesis and increased protein synthesis, synthesis and assembly of contractile proteins, and mitochondrial biogenesis. We hypothesized that CHOP would impair muscle growth in both male and female juvenile mice and that RNA sequencing would reveal disruptions in these key pathways, providing mechanistic targets for future therapeutic interventions.

## Methods

### Animals

All methods were approved by Baylor University’s Institutional Animal Care and Use Committee at protocol #2146454. All animals were kept on a 12:12 h light-dark cycle with ad libitum access to food and water. C57Bl/6 mice were bred; pups were weaned at postnatal (P) day 21 (P21) and kept with littermates of the same sex. Cages were supplemented with hydrogel (Clear H^2^O Inc, Westbrook, ME) for hydration. At P28, animals were divided into PBS or CHOP treatment. This created four groups: Male PBS (n=11), Male CHOP (n=12), Female PBS (N=9) and Female CHOP (n=8).

### CHOP Chemotherapy

The recommended chemotherapy regimen for adult NHL patients typically comprises CHOP administered every 21 days for 6–8 cycles (11), whereas pediatric lymphoma treatment typically modifies and/or intensifies the regimen based on risk stratification. Our approach was based on an adult mouse model where two cycles of CHOP effectively reduced A20 lymphoma cancer cell volume to near zero (12) and dosing modified for pediatric mice (13). Each chemotherapy cycle was initiated at P28, was 10 days long, and administered as follows: Day 1: Animals received two sequential intraperitoneal (IP) injections. The first injection contained cyclophosphamide (40 mg/kg), doxorubicin (3.3 mg/kg), and vincristine (0.05 mg/kg). Following an ∼10-minute interval, a second injection of prednisone (0.1 mg/kg). Days 2–5: Animals received one daily IP injection of prednisone (0.1 mg/kg). The CHOP regimen was followed by 5 recovery days. Animals underwent two complete cycles (20 days). Food pellets were placed on cage floor and supplemented with Nutra-Gel Diet^™^ (Bio-Serv Product #S5769) and hydrogel on treatment days to improve access to nutrition. Three male CHOP and one female CHOP mouse died following chemotherapy treatment. Tissues were collected at P48 (a study design schematic is shown in Figure 1A).

**Figure 1.**
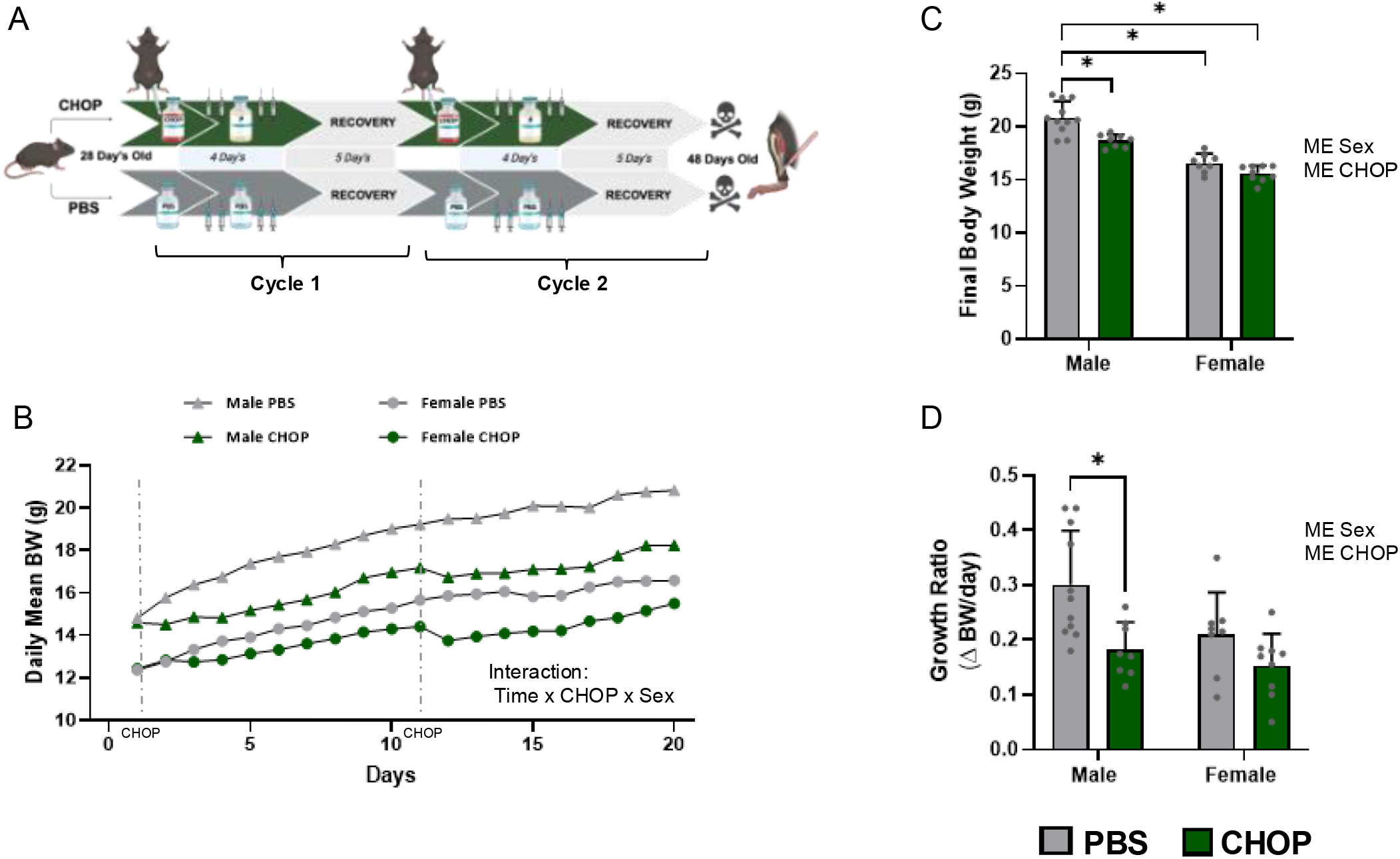
Changes in body weight following two cycles of CHOP chemotherapy in male and female mice. **A)** Study timeline showing P28 male and female C57Bl/6 mice received either PBS vehicle (grey) or CHOP chemotherapy (green) for two cycles where each cycle consisted of five days of dosing followed by five days recovery and tissues were collected a P48. **B)** Daily mean body weight across the 20-day study demonstrated expected growth in all groups, with a significant interaction of Time × CHOP × Sex. **C)** CHOP treatment and sex each had significant main effects (ME); pairwise differences demonstrated higher body weight in Male PBS compared to all other groups, indicated CHOP led to lower body weight in males, but there was no difference in body weight in females. D) Growth ratio (ΔBW/day) calculated as (final body weight – baseline body weight)/study days, showed CHOP blunted body growth in males but not females. Data are mean ± SD; main effects (ME) and interactions are annotated next to panels, ^*^p < 0.05.

### Immunohistochemistry (IHC)

IHC methods are described in detail in supplemental file 1 and previously described (14). Briefly, gastrocnemius muscles were sectioned and stained for dystrophin and myosin heavy-chain isoforms, with nuclei counterstained by DAPI. Images were acquired on a Zeiss AxioImager M2 and analyzed with MyoVision software to quantify fiber-type cross-sectional area. For Pax7^+^ Cell Detection, gastrocnemius sections were fixed in 4% PFA, heat-retrieved in citrate buffer, and blocked in 2% BSA/M.O.M. Sections were then incubated with anti-Pax7 and anti-laminin primary antibodies, followed by a biotinylated secondary, streptavidin-HRP, and TSA-AF568 amplification, with DAPI counterstain. Imaging was performed on a Zeiss AxioImager M2, and Pax7^+^ nuclei were quantified per myofiber.

### RNAseq Data Analysis

Total RNA was isolated from gastrocnemius was performed using the Qiagen miRNeasy Mini Kit (Qiagen, Hilden, Germany) according to manufacturer’s instructions. RNA samples were shipped to Oklahoma Medical Research Foundation Clinical Genomics Center for library preparation (polyA, non-strand specific) and sequencing (20 million PE150 reads, Illumina NovaSeq X Plus 25B PE150 flow cell). RNA-Seq reads were quality-controlled with FastQC (v0.11.9), adapters were trimmed using Fastp (v0.23.4), and remaining reads were aligned to the GRCm39 genome with STAR (v2.7.9a). Resulting BAM files were indexed, filtered for unmapped or low-coverage reads using SAMtools (v1.15), and gene counts were acquired using HTSeq (v2.0.2). Differential expression between PBS and CHOP treatments was analyzed in R (v4.4.2) with DESeq2 (v1.46.0), and P-values were adjusted using the Benjamini and Hochberg false discovery rate correction (15). Genes with an adjusted P value < 0.05 found by DESeq2 were assigned as differentially expressed genes (DEGs). Gene Ontology (GO) enrichment analysis of DEGs was implemented by the enrichR R package (v3.4). REVIGO (16) was used to consolidate redundant significant GO terms and to visualize terms as previously described (17).

### Statistical Analysis

Longitudinal body weight was analyzed using three-way mixed ANOVAs with the factors of Sex, CHOP and Time. Data sets comparing two factors, CHOP and SEX, and interactions among factors were analyzed using two-way ANOVAs. Significance was set at p < 0.05. Tukey post-hoc tests were performed when significant f values were found. An independent t-test was used when comparing males to females using percent difference calculations. IBM Statistical Package for the Social Sciences 28 (IBM SPSS Statistics, Cary, NC, USA) and GraphPad Prism (La Jolla, CA, USA) were used to analyze data and generate figures. Data are expressed as mean ± standard deviation. Means and results for all statistical tests can be found in Supplemental Table 2.

## Results

### CHOP chemotherapy slows growth of juvenile mice

Daily body weights over the 20-day protocol showed that CHOP treatment significantly slowed growth compared to PBS controls (Figure 1B). Both treatment and sex influenced weight trajectories, as CHOP-treated mice gained less weight overall, and males weighed more than females. A significant three-way interaction indicates that CHOP’s impact on weight gain was greater in males than females. By P48, CHOP-treated animals had lower final body weights, with a significant difference in males but not in females (Figure 1C). Calculation of daily growth rates confirmed that CHOP lowered weight gain and that this effect was most pronounced in males (Figure 1D). Overall, these results demonstrate that pediatric CHOP impairs growth, with male mice exhibiting greater sensitivity.

### CHOP induces sex-specific blunting of muscle growth

CHOP chemotherapy blunted muscle growth in a muscle- and sex-specific manner. In the soleus, CHOP had no effect in males but resulted in lower mass in females (Figure 2A). In contrast, plantaris were markedly lower in CHOP-treated males, but not in females (Figure 2B). In the gastrocnemius, the major muscle of the lower hindlimb, CHOP led to lower muscle mass in both males and females (Figure 2C). At the fiber level, CHOP resulted in lower mean gastrocnemius fiber CSA by 20–25 percent in both sexes (Figure 2D). Type IIb/x fibers appear to drive this difference because the Type IIA fibers showed no effect of Sex or CHOP. Note that Type I fibers were not analyzed because they were, on average, <2% of the fibers. When fiber CSA was expressed relative to sex-matched PBS controls, CSA loss appeared slightly greater in males than females (p=0.08, Figure 2E). Collectively, these results demonstrate that administration of CHOP to pediatric mice impaired muscle growth, predominantly affecting fast-twitch fibers, and highlight the vulnerability of growing skeletal muscles to chemotherapy.

**Figure 2.**
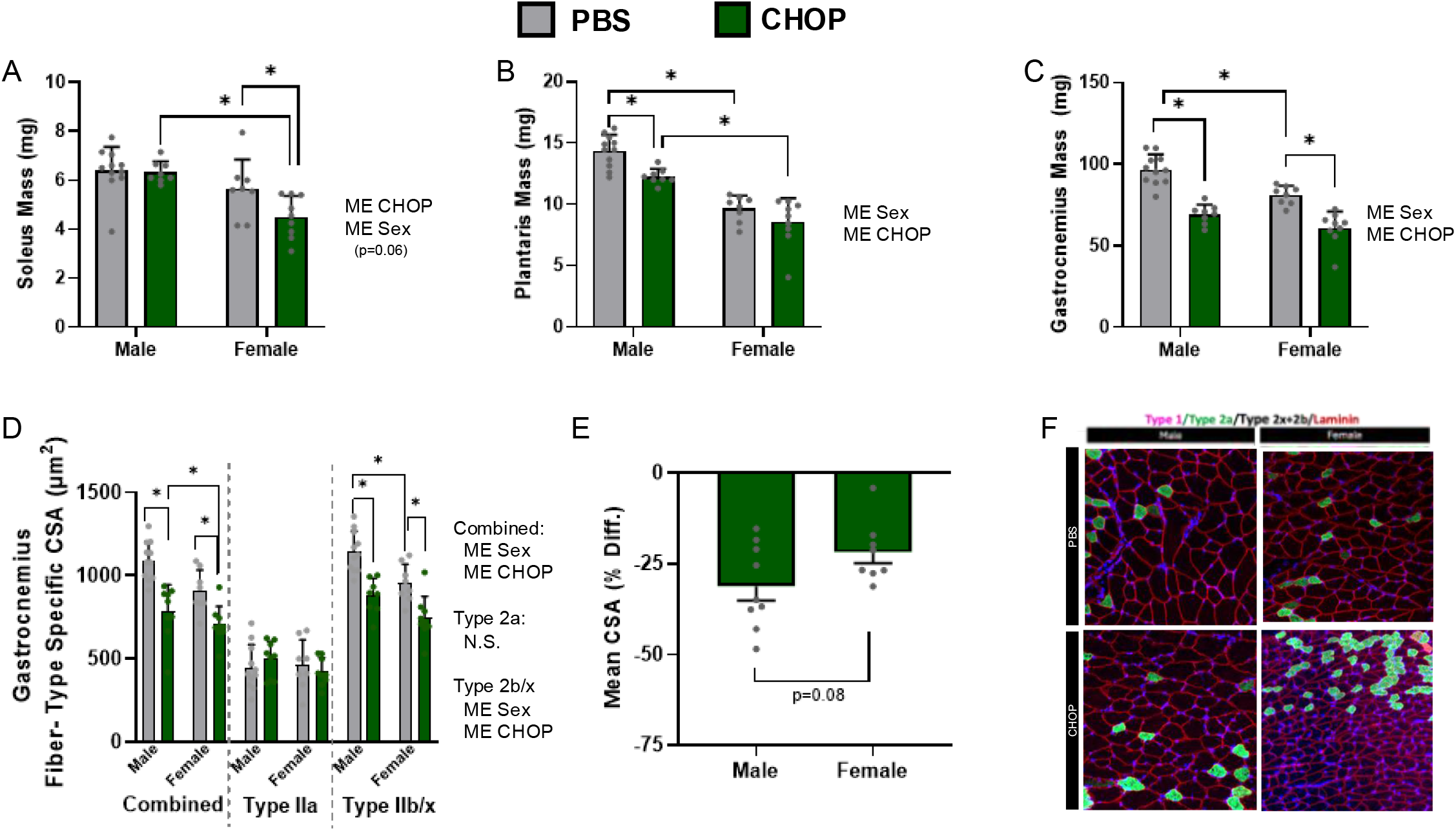
CHOP chemotherapy results in lower muscle size. **A)** CHOP led to significantly lower soleus mass, and sex also had a significant effect (ME CHOP; ME Sex), with pairwise comparison between PBS and CHOP within sex showed only an effect in females. **B)** Plantaris muscle mass was lower in CHOP-treated mice, and both treatment and sex exerted main effects on plantaris mass (ME CHOP; ME Sex), with pairwise comparison between PBS and CHOP within sex showed only an effect in males. **C)** CHOP chemotherapy blunted gastrocnemius muscle mass in both male and female mice (ME CHOP; ME Sex), with comparable atrophy observed across sexes. **D)** CHOP led to blunted combined and Type IIb/x gastrocnemius fiber cross-sectional area (CSA) (ME CHOP; ME Sex), whereas Type IIa CSA was not affected **E)** Percent difference in mean CSA relative to PBS controls showed a trend toward greater atrophy in males (p = 0.08). **F)** Representative immunofluorescence images of gastrocnemius sections stained for Type I (magenta), Type IIa (green), and laminin (red) in male and female mice treated with PBS or CHOP are shown (20×, scale bar = 50 μm). Data are mean ± SD; main effects (ME) and interactions are annotated next to panels, ^*^p < 0.05.

### RNA-Seq identified sex-specific transcriptomic responses to CHOP

To explore transcriptional changes that could underpin sex-specific differences in muscle mass, we performed bulk RNA-seq on the gastrocnemius. Males and females had a similar number of DEGs when comparing CHOP to PBS, 217 for males and 214 for females, respectively (Figure 3A-E). Despite the similar number of DEGs, there was very little overlap between the transcripts (Figure 3D), as there were only 29 shared DEGs (20 up and 9 down). Males had 188 unique DEGs, while 185 were unique in females. We have highlighted some genes within each sex that have well established roles in muscle growth and development (Figure 3A). In males, CHOP treatment results in lower expression in markers of satellite cell biology, such as Calcr, *Myf5 and Myod1*, and higher expression of *EIF2AK2*, a negative regulator of protein synthesis. In females, higher expression of Postn, which is involved in tissue remodeling and has been linked to muscle damage and fibrosis in atrophic conditions (18). There was greater expression of upstream regulators of proteolytic genes, specifically Eef1a1, Ube2c, and Ube2b, which suggests altered protein turnover. Additionally, there was higher expression of cell cycle regulators, Cdkn1a, Cdkn1c, Cdk1, and Ccna2. Of the 29 genes with concordant responses in both sexes, CHOP treatment increased expression of Mrc1, a marker of macrophage infiltration, as well as the cell-cycle regulators Cirpil and Mki67.

**Figure 3.**
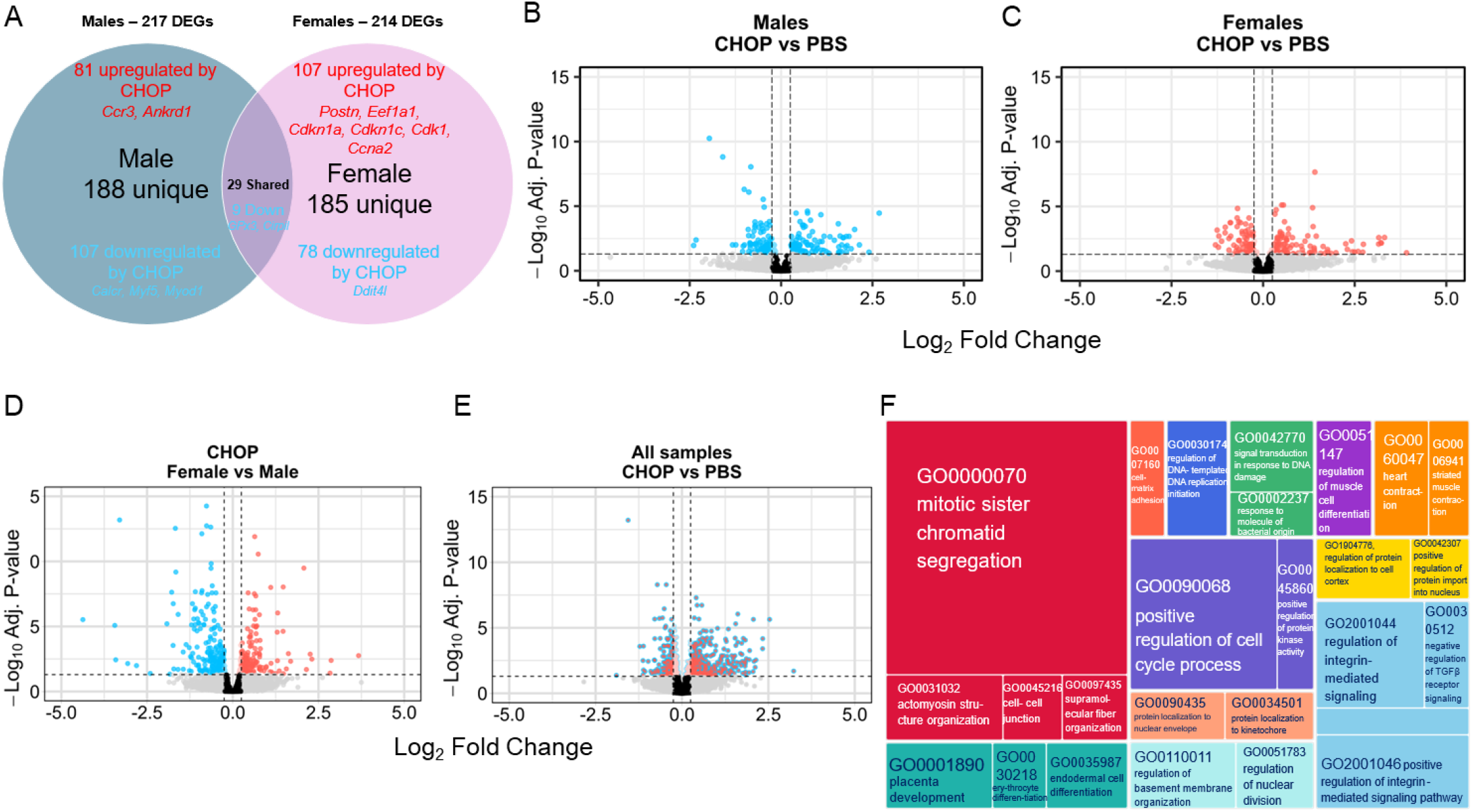
CHOP chemotherapy elicits sex-specific transcriptomic changes in gastrocnemius muscle. **A)** The Venn diagram illustrates that CHOP treatment significantly upregulated 81 genes and downregulated 107 genes uniquely in males (blue circle), upregulated 107 genes and downregulated 78 genes uniquely in females (pink circle), and altered 29 genes in both sexes (overlap) (FDR < 0.05). **C)** Volcano plot of female CHOP versus PBS comparisons shows 107 upregulated (red) and 78 downregulated genes using the same significance thresholds as in B. **D)** Volcano plot comparing CHOP effects between females and males highlights sex-biased differential expression in response to chemotherapy. **E)** Combined analysis of all samples (CHOP vs PBS) reveals the overall transcriptomic impact of CHOP across sexes. **F)** Treemap of enriched Gene Ontology biological processes among CHOP-regulated genes shows that “mitotic sister chromatid segregation” (GO:0000070) and “positive regulation of cell cycle process” (GO:0090068) are the largest terms; GO terms were summarized with Revigo, with box size proportional to term frequency and color indicating GO category.

We then pooled CHOP-treated samples and compared them to PBS controls (Figure 3 E) to identify DEGs and subjected these to GO analysis. Consolidation of significant GO Biological Process terms (Figure 3F) revealed enrichment of processes related to cell-cycle, cytoskeletal and extracellular-matrix remodeling, and regulation of muscle differentiation.

### CHOP lowers satellite cells and markers of satellite cell dynamics

Because mature myofibers are terminally differentiated, detecting a strong cell-cycle and mitotic signature in CHOP-treated muscle, especially given that treatments were administered during ongoing postnatal growth, implicates a potential role of satellite cells. To visualize how these sex-specific responses map onto distinct biological programs, we used a chord diagram to link differentially expressed genes in males and females to three key GO categories: cytoplasmic translation, ER-stress response, and muscle-cell-fate commitment (Figure 4A).

**Figure 4.**
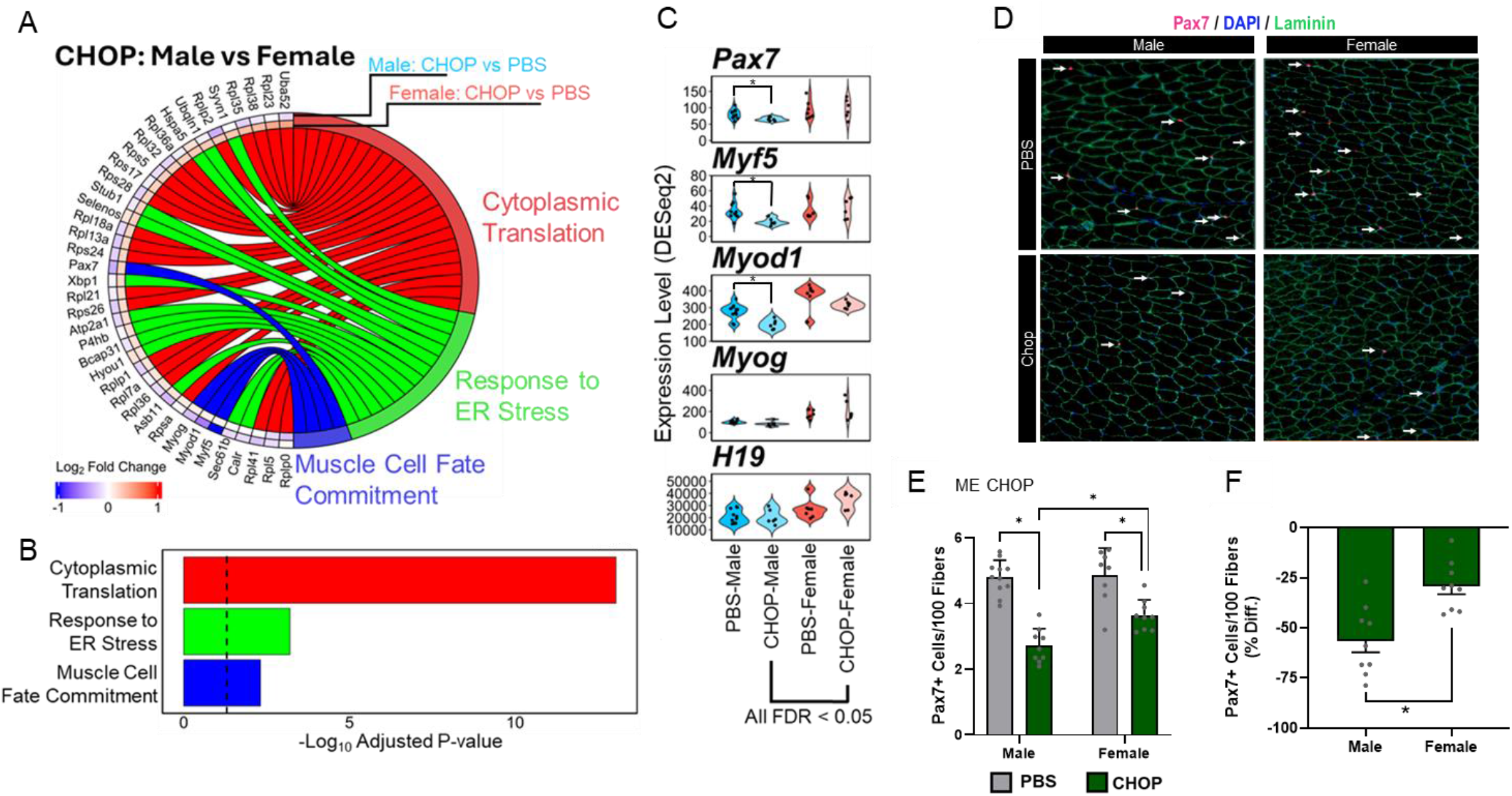
CHOP alters satellite cell commitment networks and results in less Pax7^+^ satellite cells in male and female mice. **A)** Achord diagram of log_²_ fold-changes for CHOP versus PBS in plantaris muscle highlights genes within cytoplasmic translation (red), ER stress response (green), and muscle cell fate commitment (blue); male changes are shown in blue and female changes in red (FDR < 0.05). **B)** GO-term enrichment for CHOP-regulated genes shows that cytoplasmic translation was most significantly enriched, followed by ER stress response and muscle cell fate commitment; **C)** Violin plots of DESeq2-normalized counts for Pax7, Myf5, Myod1, Myog and the long non- coding RNA H19 in PBS and CHOP groups. CHOP treated males had significantly lower Pax7, Myf5, Myod1 compared to Male PBS and were lower than Female CHOP in all transcripts **D)** Representative immunofluorescence images of gastrocnemius cross-sections stained for Pax7^+^ (pink), DAPI (blue) and laminin (green); white arrows indicate Pax7^+^ satellite cells (20×, scale bar = 50 μm). **E)** Quantification of Pax7^+^ cells per 100 fibers in gastrocnemius sections reveals less satellites cells present with CHOP in both sexes (ME CHOP) and when expressed as **F)** percent difference, data demonstrate that CHOP induced a greater relative difference of satellite cells in males compared to females. Data are mean ± SD; main effects (ME) and interactions are annotated next to panels.

CHOP treatment elicited divergent transcriptional programs in gastrocnemius muscle. Females showed a pronounced upregulation of ribosomal and translation machinery genes, whereas both sexes induced ER-stress markers. In contrast, males exhibited selective downregulation of myogenic commitment factors such as *Pax7* and Myod1. GO enrichment confirmed these patterns, with the strongest significance for translation processes, moderate enrichment for ER stress, and a notable depletion of myogenic commitment pathways (Figure 4B). Together, these data indicate that CHOP broadly perturbs protein homeostasis in developing muscle and preferentially disrupts satellite-cell–mediated differentiation in males. To further explore this possibility, we selected five well established genes from the RNAseq data that sequentially control satellite cells from quiescence to fusion (Figure 4C). Males had lower Pax7, Myf5, and MyoD. When comparing male and female CHOP, males had lower expression in all myogenic progression transcripts Pax7, Myf5, Myod1, Myog, including the lncRNA, H19, a marker of newly fused myonuclei (19).

To determine if the transcriptomic changes may have had a physiological effect, we measured the number of satellite cells (Pax7^+^) in gastrocnemius cross-sections (Figure 4D,E). There were MEs of CHOP and Sex, along with a CHOP x Sex interaction. Post-hoc comparisons revealed that CHOP led to lower Pax7^+^ cell abundance in both male and female mice (p < 0.001). Notably, CHOP-treated males exhibited significantly fewer Pax7^+^ cells compared to their female counterparts. The greater reduction in satellite-cell content in males becomes apparent when expressing CHOP-treated values as a percentage of PBS controls: males exhibited ∼60% fewer satellite cells, whereas females exhibited ∼40% fewer (Figure 4 F). Collectively, these findings indicate that CHOP disrupts satellite-cell dynamics in juvenile mice of both sexes, with a more pronounced effect in male muscle.

## Discussion

As survivorship in pediatric and AYA cancers now exceeds 85%, acute chemotherapy-induced impairments to muscle growth during adolescence, a period when muscle mass and quality critically shape lifelong metabolic and functional health, may predispose survivors to lasting weakness, metabolic dysfunction, and premature frailty (20, 21). In this study, we provide novel data demonstrating that administration of a CHOP regimen administered over two 10-day cycles in P28 mice, slows overall growth and blunts muscle CSA size in both sexes, with males exhibiting a larger deficit. Transcriptomic profiling revealed largely nonoverlapping gene-expression changes in CHOP-treated males versus females, and GO analysis pointed to satellite cell-related pathways, a finding confirmed by fewer Pax7^+^ cells, preferentially in male mice.

CHOP achieves high efficacy without radiotherapy in pediatric non-Hodgkin lymphoma by leveraging multiple agents with complementary mechanisms, including DNA crosslinking by cyclophosphamide, anthracycline-induced oxidative stress from doxorubicin, microtubule destabilization via vincristine, and glucocorticoid effects of prednisone (22). Each of these agents are considered among the five most essential childhood cancer medicines because they are incorporated into other multi-drug pediatric chemotherapy protocols (23, 24). Despite the clinical relevance, off-target effects on muscle in pediatric models remain sparse. For example, juvenile mice treated with doxorubicin exhibit ∼15-20 percent reduction in gastrocnemius CSA (5), ∼20% in soleus muscle of mice treated with vincristine (25), and a FOLFIRI regimen produced ∼25 percent lower gastrocnemius mass (10), all in male mice. These data are consistent with our model, as we observed 20-25% smaller gastrocnemius muscle fibers following CHOP. Overall, our pediatric model of CHOP chemotherapy aligns with the few other pediatric chemotherapy models and the large body of literature showing negative effects on muscle size in adult models (26). Because data on pediatric combination regimens are limited, it remains unclear whether multi-agent therapies worsen muscle wasting compared to single agents. Our results provide novel data on the acute effects of a clinically relevant pediatric CHOP regimen.

Although CHOP acutely impaired muscle growth in both male and female mice, the magnitude and molecular signature of these effects differed by sex. CHOP-treated males exhibited greater deficits in body weight gain and percent difference in gastrocnemius muscle mass than females, and transcriptomic profiling revealed largely nonoverlapping sets of DEGs in each sex. Such sexual dimorphism is not unique to our pediatric model. In adult cancer patients, Stephens et al. demonstrated that males lose more lower-limb muscle mass during cachexia than females (27), and preclinical studies similarly report sex-dependent atrophy and stem-cell responses to both tumors (28) and chemotherapy (29-31). Together, these observations underscore biological sex as a key determinant of muscle responses to CHOP.

Our RNA-Seq analysis revealed largely nonoverlapping gene expression changes in male and female gastrocnemius following CHOP treatment, with each sex engaging distinct biological programs. In males, CHOP induced downregulation of myogenic commitment factors (*Pax7, Myf5, Myod1*) and upregulation of the negative translational regulator *Eif2ak2*, consistent with impaired progenitor maintenance and suppressed protein synthesis. In contrast, females showed pronounced upregulation of extracellular-matrix and remodeling genes (e.g., Postn), increased expression of translational repressors (*Eif4ebp1*) and proteolytic pre-regulators (*Eef1a1, Ube2c, Ube2b*), and induction of cell-cycle regulators (*Cdkn1a, Cdkn1c, Cdk1, Ccna2*). Both sexes activated ER-stress markers, but only females robustly engaged ribosomal and translation machinery genes. These divergent patterns underscore a fundamental sexual dimorphism in the muscle’s molecular response to CHOP.

In our pediatric CHOP model, GO enrichment of cell-division/cell-cycle biological processes, together with differential expression of satellite-cell markers and reduced Pax7^+^ cell counts, indicates a clear disruption of muscle satellite cell dynamics. In adult chemotherapy models, however, transcriptomic signatures diverge markedly. In male mice treated with doxorubicin, p53-related genes were upregulated, implicating mitochondrial quality control dysregulation and pro-apoptotic signaling, while *Ki67* and *Ddit4* remained unchanged and no satellite-cell– associated pathways were enriched (32). There were a few similarities in that *Myod1* expression was lower and extracellular-matrix organization pathways were implicated (32). Similarly, in an adult FOLFIRI model, muscle transcripts exhibited downregulation of mitochondrial and lipid-metabolism genes and paradoxical upregulation of *Fhl3* and *Pax3*, changes inconsistent with impaired satellite-cell proliferation or differentiation (33). Although both adult studies implicate mitochondrial dysfunction and autophagy pathways in driving muscle atrophy, these signatures were not evident in our pediatric CHOP model. Moreover, neither adult study captured the pronounced satellite cell perturbations characteristic of our pediatric CHOP treatment. These contrasting molecular profiles underscore the dichotomy between adult and rapidly growing mice to chemotherapy.

The myonuclear domain, defined the cytoplasmic volume governed by each nucleus, determines a fiber’s capacity for transcription and translation (34). During postnatal growth, addition of new fibers and longitudinal growth of existing fibers rely on satellite cells to donate new myonuclei and preserve this domain. Satellite cells activate, proliferate, and fuse with growing myofibers under the control of Myf5 and MyoD1, then differentiate via Myogenin to expand fiber size. Ablation of Pax7^+^ cells by P10 produces smaller fiber CSA and reduces survival (35-37). Lineage-tracing studies show that satellite cell-derived nuclei accrue through P42 in mice (38-40) and into adolescence in rats (41, 42), establishing their ongoing role in juvenile muscle maturation.

Chemotherapeutics preferentially target proliferating cells and inhibit myoblast proliferation in vitro (43). Our CHOP regimen, administered from P28 to P48, overlaps the postnatal window of active myonuclear accretion and resulted in a diminished pool of Pax7^+^ satellite cells of ∼60 percent in males and ∼40 percent in females relative to PBS controls. Similarly, doxorubicin reduced satellite cells by ∼50% in the soleus and 40% in the EDL of nine-week-old rats (44). In contrast, vincristine alone did not negatively impact satellite cell number in the extensor digitorum longus (EDL) or soleus muscle of juvenile mice. (25). These data indicate that CHOP-induced satellite-cell loss mirrors doxorubicin’s known cytotoxicity whereas vincristine alone does not.

Our study was designed to capture the acute effects of a pediatric CHOP regimen and therefore does not address whether the observed satellite-cell depletion and fiber-size deficits persist or recover over time. In juvenile chemoradiation models, hindlimb-targeted γ-irradiation combined with vincristine to kill tumors produces sustained reductions in muscle weight, fiber CSA, and satellite cell (45, 46). Because myonuclear content is tightly correlated with fiber CSA during prepubertal growth (47), reduced satellite cells compromise muscle maturation and contribute to the well-established long-term muscle deficits (2, 10). Future work should extend analyses into late adolescence and adulthood, incorporating functional assays and repeated satellite cell and myonuclear measurements and explore the possibility of targeting satellite cells for therapeutic intervention.

Collectively, our data demonstrate that a pediatric CHOP regimen impairs overall growth, muscle mass, and fiber size in both sexes. These phenotypic deficits are underpinned by sex-specific transcriptomic programs and a pronounced depletion of Pax7^+^ satellite cells, indicating direct disruption of the muscle stem cell niche. Given the tight correlation between myonuclear content and fiber cross-sectional area during development (47) and evidence from chemoradiation models showing persistent satellite cell loss leads to persistent deficits in muscle mass (45, 46), these alterations may have long-term consequences. Our study’s focus on acute outcomes limits insight into functional consequences and the durability of satellite cell deficits. Future work should incorporate longitudinal and muscle function assessments, in addition to interventions aimed at preserving or restoring the myogenic progenitor pool. Ultimately, strategies that preserve satellite cell dynamics during chemotherapy may prove essential for maintaining muscle health in pediatric cancer survivors.

## Glossary

AYA: Adolescent/Young Adult
CHOP: Combination chemotherapy regimen comprising Cyclophosphamide, Hydroxydaunorubicin (Doxorubicin), Oncovin (Vincristine), and Prednisone
CSA: Cross-Sectional Area
DEG/DEGs: Differentially Expressed Gene(s)
DAPI: 4′,6-Diamidino-2-Phenylindole (nuclear fluorescent stain)
DSHB: Developmental Studies Hybridoma Bank
EDL: Extensor Digitorum Longus
FOLFIRI: Combination chemotherapy regimen of 5-Fluorouracil, Leucovorin, and Irinotecan
GO: Gene Ontology
IP: Intraperitoneal
IHC: Immunohistochemistry
MHC: Myosin Heavy Chain
NHL: Non-Hodgkin Lymphoma
PBS: Phosphate-Buffered Saline
PE: Paired-End (sequencing)
P: Postnatal Day
PBS: Phosphate-Buffered Saline
RNA-Seq: RNA Sequencing
REVIGO: REduce and VIsualize Gene Ontology
SAMtools: Sequence Alignment/Map tools (for processing BAM files)
SOL: Soleus (muscle)
STAR: Spliced Transcripts Alignment to a Reference (RNA-Seq aligner)

## Acknowledgments

We thank the Mooney Lab members and the Baylor vivarium staff for their support and animal care. Biorender.com was used to create Figure 1A.

## Data Management

RNA sequencing data will be deposited to NCBI Gene Expression Omnibus after pending acceptance

## Authors’ Contributions

MAM, CMD, and MPW conceived and designed the experiment. JD, MCM, MAM, CMCN, JMW, CMD, and MPW performed the experiments. JD, CMC, MA, NTT, YW, CMD, MPW analyzed data. JD, CMC, MA, NTT, YW, SVW, CMD, and MPW Interpreted results of experiments. JD, NTT, MPW prepared figures. JD and MPW drafted manuscript. All authors revised approved the final version of the manuscript.

