## Supplementary material for "Pediatric CHOP Chemotherapy Acutely Disrupts Satellite-Cell Dynamics and Blunts Muscle Mass in a Sex-Specific Manner": Supplental Tables

Supplemental File 1.

*Fiber type IHC*

Fiber type cross-sectional area was determined by IHC in the gastrocnemius muscle as previously described(14). Briefly, muscles were cut into 8μm-thick sections, and slides were incubated overnight at 4°C in primary antibodies against dystrophin and myosin heavy chain (MHC) Type 1 Developmental Studied Hybridoma Bank [DSHB]) in myosin heavy chain Type 2a . On the following day, slides were washed in PBS and incubated for 90 min at room temperature in goat anti-mouse IgG1 AF488 secondary antibody, goat anti-mouse IgG2b AF647 secondary antibody, and goat anti-rabbit IgG AF568 secondary antibody in PBS. Slides were then incubated in DAPI for 10 min followed by another wash in PBS. Slides were mounted using PBS and glycerol (1:1). A Zeiss upright microscope (AxioImager M2; Zeiss, Oberkochen, Germany) was used to capture images of the whole mounted gastrocnemius. MyoVision software was used for automated analysis of myonuclear density normalized to myofiber number, fiber type distribution, and fiber type-specific CSA (15). Unstained myofibers were counted as MHC IIx + IIb fibers (Type IIb/x).

*Detection of Pax7+ cells*

Sections were initially fixed in 4% PFA for 10 min, rinsed in PBS, treated with 3% $H_2O_2$ for 10 min, and rinsed again in PBS. Heat-mediated antigen retrieval was carried out in 10 mM sodium citrate (pH 6.5): sections were immersed in 10 mM sodium citrate at 65 °C and then gradually brought to 92 °C over ~20 min in a water bath. Once at 92 °C, they remained in the citrate buffer for 12 min before cooling for 60 min. After a PBS wash, sections were blocked for 60 min in 2% BSA with M.O.M. (Vector Labs), washed in PBS, and incubated overnight in primary antibodies against Pax7 and laminin diluted in 2% BSA. The next day, sections were washed in PBS and incubated for 90 min in Ms IgG1 Biotin secondary antibody in 2% BSA, washed again, then incubated for 75 min in streptavidin-HRP and Rb IgG AF488 diluted in PBS. Following a final PBS wash, sections were incubated for 10 min in TSA AF568 prepared in DAPI staining solution, washed in PBS, and mounted in a 1:1 PBS:Glycerol mixture under coverslips.

Table 1 – IHC antibodies

| | **Antibody** | **Source** | **Concentration** | **Conditions** |
|---|---|---|---|---|
| **Primary Antibodies** | | | | |
| | Dystrophin (rabbit IgG) | Abcam 15277 | 1:100 | 4°C overnight |
| | MHC Type 1 (mouse IgG2b) | Developmental Studied Hybridoma Bank BA.D5 | 1:100 | 4°C overnight |
| | myosin heavy chain Type 2a (mouse IgG1) | Developmental Studied Hybridoma Bank SC.71 | 1:100 | 4°C overnight |
| | PAX7 (mouse IgG1) | Developmental Studies Hybridoma Bank | 1:100 | 4°C overnight |

| | | | | |
|---|---|---|---|---|
| | Laminin (rabbit IgG) | Millipore Sigma L9393 | 1:200 | 4°C overnight |
| **Secondary Antibodies and Staining Solutions** | | | | |
| | goat anti-mouse IgG1 AF488 | Invitrogen #A21121 | 1:200 | Room Temp, 90 min |
| | goat anti-mouse IgG2b AF647 | Invitrogen #A21242 | 1:200 | Room Temp, 90 min |
| | goat anti-rabbit IgG AF568 | Invitrogen #A11011 | 1:200 | Room Temp, 90 min |
| | Mouse IgG1 Biotin | Jackson ImmunoResearch 115-065-205 | 1:1000 | Room Temp, 90 min |
| | streptavidin-HRP | Invitrogen S-911 | 1:500 | Room Temp, 75 min |
| | Rb IgG AF488 | Invitrogen #A11008, | 1:200 | Room Temp, 90 min |
| | TSA AF568 | Invitrogen #B40956 | 1:500 | Room Temp, 10 min |
| | DAPI staining solution | Invitrogen #D1306 | 10nM | Room Temp, 10 min |

**Table 2. Means, Standard Deviations, and Results of Statistical Tests**

| Variable | Male PBS | Male CHOP | Female PBS | Female Chop | ME CHOP | ME Sex | Interaction | Posthoc within CHOP | | Posthoc within Sex | |
|---|---|---|---|---|---|---|---|---|---|---|---|
| | | | | | | | | PBS M vs F | CHOP M vs F | Male PBS vs CHOP | Female PBS vs CHOP |
| Final body weight (Fig 1C) | 20.85 ±1.44 | 18.67 ±0.57 | 16.60 ±0.84 | 15.61 ±0.71 | p = 0.001 | p <0.001 | p = 0.09 | p < 0.001 | p < 0.001 | p < 0.001 | p = 0.248 |
| Growth Rate (Fig 1D) | 0.300 ±0.09 | 0.182 ±0.18 | 0.211 ±0.21 | 0.153 ±0.15 | p = 0.002 | p = 0.026 | p = 0.236 | p = 0.060 | p = 0.903 | p = 0.008 | p =0.042 |
| Soleus Mass (Fig 2A) | 6.39 ±0.93 | 6.33 ±0.41 | 5.64 ±1.13 | 4.51 ±0.81 | p < 0.001 | p = 0.061 | p = 0.09 | p = 0.088 | p < 0.001 | p = 0.886 | p = 0.016 |
| Plantaris Mass (Fig 2B) | 14.35 ±1.25 | 12.28 ±0.59 | 9.66 ±0.99 | 8.56 ±1.85 | p = 0.0014 | p <0.0001 | p = 0.290 | p < 0.001 | p < 0.001 | p = 0.002 | p = 0.104 |
| Gastroc Mass (Fig 2C) | 96.56 ±8.88 | 80.96 ±5.36 | 69.23 ±5.61 | 60.78 ±9.62 | p <0.001 | p < 0.001 | p = 0.205 | p < 0.001 | p < 0.001 | p = 0.043 | p < 0.001 |
| Gastroc Combined CSA (Fig 3D) | 1092.918 ±108.33 | 783.625 ±783.63 | 906.513 ±906.51 | 730.513 ±730.51 | p < 0.001 | p < 0.001 | p = 0.115 | p = 0.003 | p = 0.395 | p <0.001 | p = 0.006 |
| Gastroc Type IIA CSA (Fig 3D) | 592.418 ±119.18 | 491.325 ±491.33 | 573.838 ±573.84 | 450.100 ±450.10 | p = 0.404 | p = 0.009 | p = .668 | p = 0.308 | p =0.143 | p = 0.997 | p = 0.855 |
| Gastroc Type IIb/x CSA (Fig 3D) | 1146.800 ±114.13 | 831.500 ±831.50 | 954.038 ±954.04 | 764.713 ±764.71 | p < 0.001 | p < 0.001 | p = 0.157 | p = 0.003 | p = 0.395 | p <0.001 | p = 0.006 |
| Pax7+ cells/100 fibers (Fig 4E) | 2.719 ±0.52 | 4.749 ±4.75 | 3.705 ±3.70 | 2.719 ±0.52 | p <0.001 | p = 0.016 | p = 0.0321 | p = 0.999 | p = 0.008 | p < 0.001 | p < 0.001 |

| Fig. 4C Transcript | Comp-arison | Direction | Stat | Comp-arison | Direction | Stat | Comp-arison | Direction | Stat |
|---|---|---|---|---|---|---|---|---|---|
| Pax7 | Male PBS vs CHOP | Lower in CHOP | p = 0.02163 | Female PBS vs CHOP | n.d. | p = 0.693942 | Male CHOP vs Female CHOP | Lower in Males | p = 0.011 |
| Myf5 | Male PBS vs CHOP | Lower in CHOP | p = 0.00012 | Female PBS vs CHOP | n.d. | p = 0.7214 | Male CHOP vs Female CHOP | Lower in Males | p = 0.007 |

| | | | | | | | | | |
|---|---|---|---|---|---|---|---|---|---|
| MyoD | Male PBS vs CHOP | Lower in CHOP | p < 0.0001 | Female PBS vs CHOP | Lower in CHOP | p = 0.012691 | Male CHOP vs Female CHOP | Lower in Males | p < 0.001 |
| Myog | Male PBS vs CHOP | Lower in CHOP | p = 0.0848 | Female PBS vs CHOP | n.d. | p = 0.546139 | Male CHOP vs Female CHOP | Lower in Males | p = 0.008 |
| H19 | Male PBS vs CHOP | n.d. | p = 0.9927 | Female PBS vs CHOP | n.d. | p = 0.101096 | Male CHOP vs Female CHOP | Lower in Males | p = 0.003 |

Table 3.z

| | log2FoldChange | padj |
|---|---|---|
| Lyz1 | -1.96109 | 5.64849233352363e-11 |
| 1700001O22Rik | -1.59556 | 1.55451670770611e-09 |
| Nmrk2 | -0.82777 | 9.0686567103639e-09 |
| Actr3b | -1.00856 | 4.99958180857563e-07 |
| Amd1 | -0.88122 | 8.17983061625464e-07 |
| Cd24a | -0.48882 | 2.94511231921122e-06 |
| Asb10 | -0.46257 | 1.18932494642984e-05 |
| Kit | 0.722004 | 2.4527172415282e-05 |
| Igf2 | 0.726312 | 3.46035205303244e-05 |
| Pimreg | 2.684697 | 3.46035205303244e-05 |
| Tubb4b | 0.410847 | 3.46035205303244e-05 |
| Apln | 1.089475 | 4.58882609040846e-05 |
| Stc2 | 0.806701 | 0.000122136 |
| Rab12 | -0.31743 | 0.000156112 |
| Tceal5 | 0.51734 | 0.000156112 |
| Slc30a2 | -0.71328 | 0.000166592 |
| Pla2g7 | -0.56149 | 0.00020044 |
| Kifc1 | 1.566473 | 0.000232844 |
| Nkain1 | -0.39403 | 0.000232844 |
| Peg3 | 0.613703 | 0.000290293 |
| Mipep | -0.30518 | 0.0002947 |
| Cntfr | -0.52206 | 0.000297692 |
| Gramd1b | 0.587313 | 0.000297692 |
| Gpx3 | -0.70845 | 0.000302511 |
| Vash1 | 0.662615 | 0.000366556 |
| Mchr1 | -0.71636 | 0.000413054 |
| Atcay | 0.61314 | 0.00044051 |
| Cbr2 | -0.60873 | 0.000446041 |
| Gm52321 | -0.38581 | 0.000496759 |
| Msrb1 | -0.50935 | 0.000572433 |
| Cacna2d4 | -0.89923 | 0.000637555 |
| Pclaf | 2.012087 | 0.000637555 |
| H2-Aa | -0.73988 | 0.00084541 |

| | | |
|---|---|---|
| Kcng4 | -0.48326 | 0.00089777 |
| Tceal7 | 0.859871 | 0.000939017 |
| Cdk19 | -0.55756 | 0.001080519 |
| Inha | 0.488686 | 0.001350459 |
| Rtkn2 | -0.88711 | 0.001898768 |
| Ccdc141 | 0.773858 | 0.002038017 |
| Cirpil | -0.77741 | 0.002053526 |
| Plk1 | 1.615675 | 0.002053526 |
| Mycn | 1.175238 | 0.002131681 |
| Ubl7 | -0.34807 | 0.002131681 |
| Gprc5b | 0.283688 | 0.002192675 |
| Slc39a12 | 1.858216 | 0.002192675 |
| Ube2c | 1.786533 | 0.002203821 |
| Kif23 | 1.337015 | 0.002323643 |
| Col4a2 | 0.457432 | 0.002377516 |
| Brca1 | 1.703067 | 0.002656322 |
| Gm16170 | 0.507215 | 0.002817363 |
| AW551984 | 1.700328 | 0.00282248 |
| 4833439L19Rik | -0.2484 | 0.003267871 |
| Oxa1l | -0.27538 | 0.003293421 |
| Myod1 | -0.48253 | 0.004036411 |
| TrnQ | -2.32573 | 0.004118494 |
| Nrp2 | 0.41026 | 0.004222646 |
| Papln | 0.59846 | 0.004222646 |
| Prc1 | 0.920664 | 0.004222646 |
| Tiam1 | -0.62569 | 0.004222646 |
| Cdca2 | 1.458353 | 0.004347029 |
| Dym | -0.30631 | 0.004347029 |
| Espl1 | 1.122343 | 0.005468859 |
| D17H6S56E-5 | 1.061857 | 0.005490904 |
| Rcc2 | 0.309323 | 0.005632051 |
| Slc40a1 | -0.4009 | 0.00616803 |
| Rian | 0.701389 | 0.006211745 |
| Cilp | 0.664477 | 0.006260956 |
| Lrrc30 | -0.57802 | 0.006270594 |
| Retnla | -0.85361 | 0.006917415 |

| | | |
|---|---|---|
| Bcl2 | -0.69351 | 0.007540588 |
| H2bc6 | 0.435414 | 0.007707508 |
| Slc15a5 | -0.83788 | 0.007707508 |
| Tubb6 | 0.370826 | 0.007827398 |
| Ccl21a | -0.82593 | 0.008265437 |
| Sqor | -0.30331 | 0.008265437 |
| Gstm1 | -0.37501 | 0.008284103 |
| Gm17224 | 0.397847 | 0.008673293 |
| S100a1 | -0.72066 | 0.0089189 |
| Psap | -0.20523 | 0.009159638 |
| Gm57536 | 0.515579 | 0.009617048 |
| Bcl6b | 0.737636 | 0.009927031 |
| Calcr | -1.32724 | 0.009927031 |
| Cenpe | 1.659935 | 0.009927031 |
| Cpt2 | -0.29911 | 0.009927031 |
| Gm53882 | -0.8423 | 0.009927031 |
| Gnpat | -0.19171 | 0.009927031 |
| Hmga2-ps1 | -1.23823 | 0.009927031 |
| Kcmf1 | -0.21444 | 0.009927031 |
| Lamp1 | -0.19459 | 0.009927031 |
| Lrrfip1 | 0.272586 | 0.009927031 |
| Med12l | 0.986287 | 0.009927031 |
| Pcdh1 | 0.339832 | 0.009927031 |
| Plaat3 | -0.21482 | 0.009927031 |
| Plxna3 | 0.815366 | 0.009927031 |
| Sema4d | -0.23037 | 0.009927031 |
| Serinc3 | -0.28542 | 0.009927031 |
| Tbc1d4 | 0.380514 | 0.009927031 |
| Troap | 2.143187 | 0.009927031 |
| 8430426J06Rik | -1.00131 | 0.010027355 |
| Fth1 | -0.40276 | 0.010027355 |
| Pcdh17 | 0.724928 | 0.010027355 |
| Pigbos1 | -0.56733 | 0.010027355 |
| Ryr3 | -0.68529 | 0.010027355 |
| Ano10 | -0.25826 | 0.010329535 |
| Mypop | -0.37991 | 0.010329535 |

| | | |
|---|---|---|
| Rnf10 | -0.1828 | 0.010329535 |
| Itga9 | 0.380202 | 0.010581039 |
| Mcm6 | 0.515008 | 0.010581039 |
| Gm53757 | -2.39144 | 0.011039762 |
| Atp1b4 | 0.838407 | 0.011104037 |
| Ckap2 | 1.807439 | 0.011104037 |
| Gm39469 | 0.510025 | 0.011104037 |
| Mgst1 | -0.50218 | 0.011104037 |
| Rnf126 | -0.3958 | 0.011663795 |
| Gde1 | -0.25332 | 0.012050521 |
| Gm15543 | -1.03447 | 0.01226295 |
| Serpinb6a | -0.42947 | 0.01226295 |
| Atcayos | 0.365678 | 0.012513557 |
| Ehd4 | 0.28038 | 0.012513557 |
| Fbxw5 | -0.21129 | 0.012513557 |
| Nab2 | 0.401033 | 0.012513557 |
| Rpn1 | -0.3062 | 0.012513557 |
| Clspn | 1.580127 | 0.014245055 |
| Mki67 | 1.766023 | 0.014520971 |
| Gm29773 | 0.34226 | 0.015417609 |
| Arap3 | 0.293304 | 0.015586888 |
| Myf5 | -0.84255 | 0.015797663 |
| Lrfn3 | -0.4597 | 0.016718312 |
| Itgb5 | -0.29298 | 0.016731535 |
| Ccdc85c | -0.27112 | 0.017032032 |
| Stab2 | -0.48692 | 0.017677961 |
| 5031439G07Rik | -0.29409 | 0.017834617 |
| Clca2 | 1.005281 | 0.018215225 |
| Enho | -0.53237 | 0.018511795 |
| Aurkb | 1.918538 | 0.019898905 |
| Armcx4 | 0.447724 | 0.01994267 |
| Chpf2 | 0.330922 | 0.01994267 |
| Ripk3 | 0.88532 | 0.01994267 |
| Gm42226 | 0.473801 | 0.02036481 |
| Hcn2 | -0.63342 | 0.02036481 |
| Clu | -0.46696 | 0.022070165 |

| | | |
|---|---|---|
| Ubqln4 | -0.20409 | 0.022070165 |
| Sec16b | 0.477396 | 0.022331533 |
| Kif3c | -0.44809 | 0.022456704 |
| Rab35 | -0.24893 | 0.02269391 |
| Rpl7l1 | -0.26457 | 0.02269391 |
| Nid2 | 0.423233 | 0.023030424 |
| Hycc1 | 0.336587 | 0.023035341 |
| Ogdh | -0.18188 | 0.023612577 |
| Scin | 0.528786 | 0.023979483 |
| Iqgap3 | 0.966582 | 0.024228779 |
| Cibar1 | -0.49491 | 0.024756392 |
| Clmn | -0.46817 | 0.024756392 |
| Kif11 | 1.107567 | 0.024756392 |
| Shc2 | -0.57505 | 0.024899398 |
| Rcan2 | -0.40353 | 0.024985249 |
| Gck | -0.70704 | 0.025787641 |
| Gm33319 | 1.164528 | 0.026178235 |
| Golm1 | -0.34226 | 0.026829769 |
| Eif2ak2 | 0.376286 | 0.026987978 |
| Mtss2 | -0.32632 | 0.028060912 |
| Arhgap11a | 0.606143 | 0.02887653 |
| Ndel1 | -0.20682 | 0.029880234 |
| Mrc1 | 0.431063 | 0.030451094 |
| Fam13a | 0.342297 | 0.030815921 |
| Rrm2 | 0.851815 | 0.030815921 |
| Arl4c | 0.669249 | 0.031291782 |
| Gypc | -0.31339 | 0.031662725 |
| Sln | 1.233801 | 0.031977367 |
| 2310015K22Rik | -0.3464 | 0.031993943 |
| 2210408F21Rik | -0.29341 | 0.032291066 |
| Apod | -0.54405 | 0.032291066 |
| Cd27 | -1.05161 | 0.032291066 |
| Fam234a | -0.20555 | 0.032291066 |
| H2-Ab1 | -0.64141 | 0.032291066 |
| Hmgb2 | 0.58245 | 0.032291066 |
| Irx3os | -0.88142 | 0.032291066 |

| | | |
|---|---|---|
| Kntc1 | 1.65074 | 0.032291066 |
| Mtfr1l | -0.20054 | 0.032291066 |
| Scmh1 | -0.26382 | 0.032291066 |
| Sh3glb1 | -0.15181 | 0.032291066 |
| Thap7 | -0.41276 | 0.032291066 |
| Supt3 | -0.47072 | 0.033226343 |
| Mmab | -0.2179 | 0.033493873 |
| Pttg1 | -0.38581 | 0.033493873 |
| Tet1 | 0.578577 | 0.03382967 |
| Stil | 1.789098 | 0.0338548 |
| Zfp358 | -0.29375 | 0.034472496 |
| Rgs16 | 0.943138 | 0.035217199 |
| Magix | -0.26022 | 0.035394369 |
| Alkbh6 | -0.32461 | 0.035395785 |
| Gstm5 | -0.37343 | 0.035395785 |
| Sh2d6 | 0.657321 | 0.035395785 |
| Tmem248 | -0.2376 | 0.035658776 |
| Ulbp1 | 1.574952 | 0.035658776 |
| A930016O22Rik | 0.480532 | 0.036504954 |
| Sapcd2 | 2.403805 | 0.036523318 |
| Xk | -0.43492 | 0.036758437 |
| Eif3e | -0.19626 | 0.037173379 |
| Coq10a | -0.31319 | 0.037703437 |
| Actc1dt | 1.621187 | 0.038101933 |
| Ccdc3 | -0.57406 | 0.038649497 |
| Nuf2 | 1.587155 | 0.038684435 |
| Car4 | -0.78071 | 0.039105393 |
| Grb10 | 0.323127 | 0.039480571 |
| Gm34256 | 0.512946 | 0.040358663 |
| Col4a1 | 0.474969 | 0.041638315 |
| Rilpl1 | -0.36093 | 0.041831702 |
| Cd74 | -0.81401 | 0.042033401 |
| Gm52697 | 1.122208 | 0.042623947 |
| Cpne2 | 0.396646 | 0.043581566 |
| Ndc80 | 1.307226 | 0.043581566 |
| Rnf130 | -0.17697 | 0.046063505 |

| | | |
|---|---|---|
| Lmnb2 | 0.376762 | 0.04610371 |
| Cspg4 | 0.320916 | 0.047207273 |
| Lcp2 | 0.753154 | 0.047852534 |
| Dnhd1 | 0.670736 | 0.048318063 |
| Ankrd1 | 1.337514 | 0.050134501 |
| Kcne5 | -0.73457 | 0.050250203 |

Table 4.

| Gene/transcript | log2FoldChange | padj |
|---|---|---|
| Hba-a2 | 1.350976 | 2.23413901906789e-09 |
| Dennd5a | 0.33885 | 3.3740967505997e-07 |
| St6galnac4 | 0.542765 | 1.13845357095075e-06 |
| Prkab2 | -0.37941 | 2.09517658835872e-06 |
| Cpeb1 | -0.70432 | 2.74976732075986e-06 |
| 2310015K22Rik | -0.70422 | 4.87225723111976e-06 |
| B3galnt2 | -0.47091 | 1.8429349863982e-05 |
| Mrc1 | 0.555973 | 4.49232583760859e-05 |
| Ppp1r3a | -0.29526 | 4.49232583760859e-05 |
| Asb14 | -0.29368 | 5.0271546417217e-05 |
| Nrp2 | 0.439061 | 8.7860440047279e-05 |
| Postn | 0.849976 | 0.000202 |
| Ociad2 | -0.40113 | 0.000239 |
| Alas2 | 1.281363 | 0.000249 |
| Ip6k3 | 0.500357 | 0.000296 |
| A930003A15Rik | 1.18533 | 0.000362 |
| Eif4ebp1 | 0.44791 | 0.000362 |
| Hbb-bt | 1.397991 | 0.000362 |
| Mzt1 | -0.33825 | 0.000362 |
| Tmed9 | 0.381552 | 0.000362 |
| Tubb6 | 0.496074 | 0.000362 |
| Maged1 | 0.41563 | 0.00037 |
| Kctd9 | -0.27243 | 0.000591 |
| Ano5 | -0.29028 | 0.000605 |

| | | |
|---|---|---|
| Cul7 | 0.490465 | 0.000685 |
| Eef1a1 | 0.320106 | 0.000685 |
| Ghr | -0.24572 | 0.000685 |
| Lyz1 | -1.07764 | 0.000685 |
| Ehd4 | 0.428633 | 0.000819 |
| Mical2 | -0.36971 | 0.000819 |
| Npr3 | 0.683918 | 0.000865 |
| Casr | -0.76702 | 0.00111 |
| Ntmt2 | -0.42957 | 0.00111 |
| Casq2 | 0.632574 | 0.001148 |
| Gm32652 | -0.82645 | 0.001401 |
| Hbb-bs | 1.130372 | 0.001401 |
| Knl1 | 2.403493 | 0.001401 |
| Hba-a1 | 1.097056 | 0.00144 |
| Igfbp5 | -0.5264 | 0.001742 |
| Rn7sk | 3.110141 | 0.001995 |
| Gm14325 | -0.38725 | 0.002019 |
| Acyp1 | -0.40754 | 0.002091 |
| Exd1 | -0.59084 | 0.002091 |
| Rhbdl3 | 0.41842 | 0.002091 |
| Mcm5 | 0.917486 | 0.002772 |
| Tnnt2 | 0.780457 | 0.00301 |
| Tubb5 | 0.567577 | 0.003024 |
| Ddit4l | -0.41384 | 0.003052 |
| Ttc9 | 0.601309 | 0.003628 |
| Osbpl6 | -0.2607 | 0.003865 |
| Zfp469 | 0.607837 | 0.003896 |
| Gm33543 | -1.05392 | 0.004311 |
| Agt | -0.44449 | 0.004414 |
| Cpeb2 | -0.56013 | 0.004619 |
| Gm52321 | -0.58153 | 0.004619 |
| Slc7a1 | 0.543016 | 0.004619 |
| Carnmt1 | -0.26945 | 0.004631 |
| Mir6236 | 3.016026 | 0.004829 |
| Rmdn1 | -0.30348 | 0.005231 |
| Slc44a2 | -0.16867 | 0.005884 |

| | | |
|---|---|---|
| F830016B08Rik | -0.64651 | 0.005892 |
| Mast4 | -0.3303 | 0.005892 |
| Psat1 | 1.022037 | 0.005892 |
| Pdia3 | 0.336393 | 0.006328 |
| Rnase13 | -1.17545 | 0.007895 |
| Gm15564 | 2.940876 | 0.008416 |
| Kif18b | 2.992716 | 0.008416 |
| Gm12514 | -1.22037 | 0.008857 |
| Cdkn1a | 0.544305 | 0.009071 |
| Lmnb2 | 0.63032 | 0.009627 |
| Eda2r | 0.439401 | 0.010022 |
| Arhgap5 | -0.29983 | 0.010092 |
| Cdh4 | -0.71331 | 0.010092 |
| Tceal8 | 0.380363 | 0.010092 |
| Setd7 | -0.26581 | 0.010351 |
| 6430548M08Rik | 0.500916 | 0.0107 |
| Ckap2l | 1.725561 | 0.0107 |
| Gm38650 | -1.00324 | 0.0107 |
| Pclaf | 2.104538 | 0.0107 |
| Zfp654 | -0.27655 | 0.010751 |
| Oxr1 | -0.2061 | 0.010871 |
| Zfp385a | 0.588648 | 0.010871 |
| Gpx3 | -0.58053 | 0.010895 |
| Il15 | -0.63893 | 0.01094 |
| Slc28a2 | -0.39705 | 0.01094 |
| Asb10 | -0.40914 | 0.011166 |
| Btbd8 | 0.367132 | 0.011166 |
| BC023105 | -0.54339 | 0.011205 |
| Arpc1b | 0.379054 | 0.011304 |
| Emp1 | 0.358501 | 0.011304 |
| Kifc1 | 1.73131 | 0.011304 |
| Calu | 0.268536 | 0.011357 |
| Togaram1 | -0.31632 | 0.011357 |
| Gm15179 | -0.42455 | 0.01208 |
| Galnt17 | 0.588353 | 0.012462 |
| Sln | 1.305054 | 0.012462 |

| | | |
|---|---|---|
| Ube2c | 1.283466 | 0.012462 |
| Hspa5 | 0.338776 | 0.013582 |
| Pimreg | 2.504843 | 0.013636 |
| Tceal7 | 1.158016 | 0.013636 |
| Dbnl | 0.298559 | 0.014234 |
| Sorbs2 | -0.3105 | 0.014408 |
| Gm53756 | 2.460214 | 0.014644 |
| Hmmr | 2.363008 | 0.016009 |
| Vcan | 0.493218 | 0.016111 |
| Celsr2 | -0.56274 | 0.016267 |
| Ifitm3 | 0.423575 | 0.016267 |
| Cilp | 0.50707 | 0.016669 |
| Vash2 | 1.012589 | 0.017872 |
| COX2 | -0.3985 | 0.018208 |
| AW551984 | 2.886913 | 0.018702 |
| Kif11 | 1.363419 | 0.018702 |
| Rnd1 | -0.8979 | 0.019499 |
| Slc20a1 | -0.38087 | 0.019594 |
| Slc25a36 | -0.27082 | 0.020248 |
| Lgals3 | 0.732988 | 0.020504 |
| Tecrl | 1.534809 | 0.021498 |
| Ccdc85c | -0.34495 | 0.021609 |
| Gm19417 | 0.673578 | 0.021609 |
| Grb2 | 0.153643 | 0.021609 |
| Mcm6 | 0.568772 | 0.021609 |
| Otulin | 0.400645 | 0.021609 |
| Rictor | -0.27947 | 0.021609 |
| Ifit3b | -0.6799 | 0.021649 |
| Gpx8 | 0.423262 | 0.021793 |
| Adra1a | -0.68586 | 0.021901 |
| Tmem164 | 0.491479 | 0.021901 |
| Arhgef26 | 0.938975 | 0.021938 |
| Frem2 | 0.605613 | 0.021938 |
| Hectd2 | -0.43027 | 0.021938 |
| Impa2 | 0.447586 | 0.021938 |
| Serpine1 | 0.792377 | 0.021938 |

| | | |
|---|---|---|
| Tmem43 | 0.253585 | 0.021938 |
| Pitpnm3 | -0.46743 | 0.022558 |
| Arhgap24 | 0.429688 | 0.022959 |
| Cyth3 | 0.290141 | 0.022959 |
| Rnpc3 | -0.30837 | 0.023321 |
| 1810014B01Rik | -0.37983 | 0.023972 |
| Dhdh | -0.33061 | 0.023972 |
| Cirpil | -0.75432 | 0.024116 |
| Pla2g5 | -0.97769 | 0.025275 |
| Siglec1 | 0.851218 | 0.025413 |
| Tubd1 | -0.30039 | 0.025422 |
| 2900052L18Rik | 0.541278 | 0.025627 |
| Clmn | -0.40354 | 0.026683 |
| Phaf1 | -0.26176 | 0.028205 |
| Shisa4 | 0.334825 | 0.028205 |
| Trp53inp1 | 0.495499 | 0.028205 |
| Ncapg | 2.450481 | 0.028294 |
| Nabp1 | -0.40108 | 0.028739 |
| Gm53856 | -0.74329 | 0.028755 |
| Hcls1 | 0.556362 | 0.028791 |
| Gm4861 | -0.58992 | 0.029009 |
| Gm54020 | -0.91106 | 0.029009 |
| Ccna2 | 1.371748 | 0.029103 |
| Pcdh17 | 0.633788 | 0.029103 |
| Ctsz | 0.264697 | 0.030264 |
| Lama4 | 0.320911 | 0.030264 |
| Uhrf1 | 1.367919 | 0.031691 |
| Cdk1 | 1.84829 | 0.031939 |
| Cdnf | -0.28644 | 0.031939 |
| Vim | 0.384054 | 0.031939 |
| 4833412C05Rik | 2.107534 | 0.031994 |
| Asb15 | -0.40932 | 0.031994 |
| Ckap2 | 1.632848 | 0.031994 |
| Gm32200 | -0.67074 | 0.032131 |
| Scrn3 | -0.21827 | 0.032131 |
| Myl9 | 0.516714 | 0.032237 |

| | | |
|---|---|---|
| Gm22513 | 4.828742 | 0.032547 |
| Hspb1 | 0.339161 | 0.032547 |
| Tmem127 | 0.209748 | 0.033489 |
| Rmrp | 3.685592 | 0.033956 |
| Hsp90b1 | 0.263932 | 0.034942 |
| Amd1 | -0.44877 | 0.035145 |
| Mki67 | 1.892934 | 0.035274 |
| Cdkn1c | 1.223762 | 0.035279 |
| C1qtnf3 | 1.227491 | 0.035318 |
| Gm53561 | -0.86572 | 0.035318 |
| Myl6b | 0.766591 | 0.035886 |
| Top2a | 1.840712 | 0.036702 |
| Mlxip | -0.20781 | 0.036841 |
| Umad1 | -0.30765 | 0.037886 |
| COX3 | -0.41173 | 0.038512 |
| Pgd | 0.384699 | 0.039463 |
| Dnaja4 | 0.303332 | 0.039833 |
| Prc1 | 0.76143 | 0.042554 |
| Csrp3 | 1.015125 | 0.042626 |
| Gm26917 | 2.218973 | 0.042626 |
| Nusap1 | 0.825619 | 0.043217 |
| Ifi30 | 0.450265 | 0.043262 |
| Adi1 | -0.2687 | 0.043318 |
| Mpzl1 | 0.325599 | 0.043318 |
| Cenpi | 1.739053 | 0.043514 |
| Esyt1 | 0.201006 | 0.043514 |
| Frmd4b | 0.353068 | 0.044123 |
| Best3 | -0.47211 | 0.04441 |
| Junos | -0.67269 | 0.04441 |
| Gm42226 | 0.323199 | 0.044576 |
| Msn | 0.258895 | 0.044576 |
| Lmcd1 | 0.469846 | 0.045408 |
| Tppp2 | -1.0186 | 0.045408 |
| Ube2b | -0.25781 | 0.045408 |
| Mrgprh | -0.58497 | 0.045531 |
| Ccdc85b | 0.366614 | 0.045948 |

| | | |
|---|---|---|
| Pck1 | 1.329452 | 0.045948 |
| Adk | -0.27578 | 0.047373 |
| Ctnna1 | 0.155711 | 0.047373 |
| Cyb5d1 | -0.19741 | 0.047373 |
| Iqgap1 | 0.250446 | 0.047373 |
| Nucb1 | 0.230529 | 0.047373 |
| Sulf2 | 0.354107 | 0.047373 |
| Rps4l | 0.501166 | 0.047754 |
| Acot9 | 0.287321 | 0.048804 |
| Ckap4 | 0.383192 | 0.049117 |
| ATP6 | -0.40467 | 0.050383 |
| Timp1 | 0.946938 | 0.054516 |
| Ubr2 | -0.24564 | 0.054516 |
| Depp1 | 1.274405 | 0.054562 |
| Esco1 | -0.27989 | 0.054562 |
| Osbpl1a | 0.234817 | 0.054562 |
| Sptan1 | 0.228636 | 0.054603 |
| Bcap31 | 0.220034 | 0.054826 |
